# The discovery of gene mutations making SARS-CoV-2 well adapted for humans: host-genome similarity analysis of 2594 genomes from China, the USA and Europe

**DOI:** 10.1101/2020.09.03.280727

**Authors:** Weitao Sun

## Abstract

Severe acute respiratory syndrome coronavirus 2 (SARS-CoV-2), a positive-sense single-stranded virus approximately 30 kb in length, causes the ongoing novel coronavirus disease-2019 (COVID-19). Studies confirmed significant genome differences between SARS-CoV-2 and SARS-CoV, suggesting that the distinctions in pathogenicity might be related to genomic diversity. However, the relationship between genomic differences and SARS-CoV-2 fitness has not been fully explained, especially for open reading frame (ORF)-encoded accessory proteins. RNA viruses have a high mutation rate, but how SARS-CoV-2 mutations accelerate adaptation is not clear. This study shows that the host-genome similarity (HGS) of SARS-CoV-2 is significantly higher than that of SARS-CoV, especially in the ORF6 and ORF8 genes encoding proteins antagonizing innate immunity *in vivo*. A power law relationship was discovered between the HGS of ORF3b, ORF6, and N and the expression of interferon (IFN)-sensitive response element (ISRE)-containing promoters. This finding implies that high HGS of SARS-CoV-2 genome may further inhibit IFN I synthesis and cause delayed host innate immunity. An ORF1ab mutation, 10818G>T, which occurred in virus populations with high HGS but rarely in low-HGS populations, was identified in 2594 genomes with geolocations of China, the USA and Europe. The 10818G>T caused the amino acid mutation M37F in the transmembrane protein nsp6. The results suggest that the ORF6 and ORF8 genes and the mutation M37F may play important roles in causing COVID-19. The findings demonstrate that HGS analysis is a promising way to identify important genes and mutations in adaptive strains, which may help in searching potential targets for pharmaceutical agents.

## Introduction

In December 2019, a novel coronavirus SARS-CoV-2 was reported as the cause of COVID-19. SARS-CoV-2 has a positive-sense single-stranded RNA with a length of approximately 30 kb[1]. Studies have shown that considerable genetic diversity exists between SARS-CoV-2 and SARS-CoV[1, 2]. Compared with SARS-CoV, SARS-CoV-2 appears to be more contagious and more adapted to humans[3]. The distinctions in pathogenicity and virulence might be related to genomic diversity.

RNA viruses are susceptible to genetic recombination, and viral populations may evolve improved adaptability in the process of infecting hosts. By comparing the genome similarity of the virus to the host, the adaptability of the virus to the host can be inferred. Although the genomes of viruses and hosts are quite different in general, nucleotide sequence similarities do exist. Such similarities may have three biological significances. (1) These similar fragments come from a common ancestor and remain stable over long-term evolution due to their biological significance. (2) Similar genomic fragments are coincidentally preserved in both viruses and hosts over time because of the biological benefits of the gene products. (3) When the virus interacts with the hosts, mutants are created by virus-host gene exchanges, causing genome similarities.

A growing number of studies on virus-host gene similarity have been reported. Simian virus 40 (SV40), the first animal virus to undergo complete full-sequence DNA analysis, can infect monkeys and humans and cause tumors[4]. Rosenberg et al.[5] found that some mutant SV40 viruses contained nucleic acid sequences from their host monkeys. This finding suggests that viruses can recombine with host genes to complete their own physiological processes, which makes up for a lack of function or increases virulence. Genes similar to specific fragments of the human genome in molluscum contagiosum virus (MCV) have been reported[6]. MCV is a human poxvirus and lacks the genes associated with virus-host interactions in other poxvirus species (variola virus). However, genes in MCV with high similarity to specific fragments of the human genome are also hard to find in other poxviruses. These host-like genes may provide MCV-specific strategies for coexistence with the host[6]. In other words, it is very likely that viruses use host-specific genes to perform activities related to virus-host interactions, such as evasion of the host innate immune system. When human peripheral blood DNA was used as a template for polymerase chain reaction (PCR), 5 of 6 samples could be amplified by Epstein-Barr virus (EBV)- or hepatitis C virus (HCV)-specific primers[7]. Therefore, it is speculated that some genes of the two viruses may also exist in the human genome or that the viruses may have homology with human genes. This hypothesis implies that not only can the virus have the host’s genes but also the host itself may have genes from the virus.

Selection pressure exerted by the host immune system plays an important role in shaping virus mutations. Homology between virus and host proteins indicates the presence of host gene capture. Evolution of viral genes may involve intergenome gene transfer and intragenome gene duplication[8]. By acquiring immune modulation genes from cells, viruses have evolved proteins that can regulate or inhibit the host’s immune system[9, 10]. A recent study showed that human genome evolution was shaped by viral infections[11]. In mammals, nearly 30% of the adaptive amino acid changes in the human proteome are caused by viruses, suggesting that viruses are one of the major driving factors for the evolution of mammalian and human proteomes[12]. These findings support the possibility that SARS-CoV-2 may exchange genetic information with host cells. It can be inferred that most of the traits and mechanisms retained in “coevolution” between viruses and their hosts, including genetic and mutational mechanisms, benefit at least one or both. At the molecular level of evolution, the exchange of genetic information is necessary for virus-host mutual adaptation, leading to the similarity of nucleotide sequences.

It is interesting to study the relationship between gene similarities and viral transmission/pathological ability. The single-stranded RNA of coronavirus generally encodes three categories of proteins: (1) the replication proteins open reading frame (ORF)1a and ORF1ab; (2) the structural proteins S (spike), E (envelope), M (membrane) and N (nucleocapsid); and (3) accessory proteins with unknown homologues. The structural protein genes are organized as ‘-S-E-M-N-’ in the SARS-CoV-2 genome, and accessory protein genes are distributed between S and E, M and N.

The accessory protein genes play a key role in inhibiting the innate immune response *in vivo* and are more susceptible than the other genes to species-specific mutations under the pressure of evolutionary selection. Once inside the cell, the virus immediately confronts other critical proteins known as host-restriction factors (HRFs)[13]. HRFs are proteins that recognize and block viral replication. Virus-host interactions control species specificity and viral infection ability. Under pressure from the host immune system, viruses must be able to overcome a range of constraints associated with the host species and often show evolutionary mutation selections. It is hypothesized that accessory ORFs may retain beneficial mutations to increase host-genome similarity (HGS). Identifying emerging genetic mutations in virus populations with high HGS may aid the understanding of how SARS-CoV-2 evolved adaptation to humans. To the best of our knowledge, studies on the genetic similarity between SARS-CoV-2 and the human genome have not been reported.

This study investigated the HGS of SARS-CoV-2 genes and elucidated the links between HGS and virus adaptation to humans. A power law relationship was discovered between the expression of genes with interferon (IFN)-stimulated response elements (ISREs) and HGS. ORFs with higher HGS suppressed the gene expression of ISRE-regulated genes to a greater extent. Applying HGS analysis to 2594 SARS-CoV-2 genomes from China, the USA and Europe, it was found that the ORF6 and ORF8 genes of SARS-CoV-2 had more significant HGS increments than SARS-CoV. In addition, three different sets of surviving mutations were identified in SARS-CoV-2 genomes for China, the USA and Europe. Interestingly, an ORF1ab mutation, 10818G>T, which resulted in the residue mutation M37F in the transmembrane protein nsp6, was observed in virus populations of all three regions. This mutation did not occur in strain populations with low HGS but gradually appeared in populations with high HGS. This finding provides strong evidence that SARS-CoV-2 may accelerate adaptation in humans through increasing HGS of the ORF6 and ORF8 genes and selecting the M37F mutation. However, the underlying mechanism by which these genes and mutations make SARS-CoV-2 more adapted to humans remains unclear.

## Materials and Methods

### Viral genome data

By using BLAST ORFfinder[14], 31 ORFs were detected in the RNA genome sequence (29903 nt) of SARS-CoV-2 (GenBank: MN908947.3). Only ATG was used as the ORF start codon, and nested ORFs were ignored. Among all the ORFs in the SARS-CoV-2 sequence, we selected the longest 14 as targets, whose lengths were no less than 75 nt. For genome comparison, ORFs in the SARS-CoV genome with a length of 29728 nt (GenBank: AY394850.2) were also identified. There were 19 ORFs with lengths no less than 75 nt in the SARS-CoV sequence.

The SARS-CoV-2 genomes were obtained from the GISAID database[15]. By May 20, 2020, the GISAID database (https://www.gisaid.org/) had 416 SARS-CoV-2 genomes from China, 5184 genomes from the USA and 10954 genomes from Europe. Complete and high-coverage genomes were used to ensure accurate HGS calculations. The sequences containing nucleotide names other than A, G, C and T were removed from the dataset. In total, 2594 SARS-CoV-2 genomes were used in the current study, including 200 from China, 1538 from the USA and 856 from Europe. The CDSs of the SARS-CoV-2 genome were identified by using MATLAB (https://www.mathworks.com/help/bioinfo/ref/seqshoworfs.html). The accession IDs of the genomes used in the article can be found in the Supplemental Information.

Human SARS-CoV genomes were collected from NCBI GenBank[16]. There were 25 CDSs of SARS-CoVs (full-length sequences only, with all ORF sequences, no nucleotide names other than A, G, C and T) at the time of article preparation. The accession IDs of these viral sequences can be found in the Supplemental Information.

### Host-genome similarity (HGS)

The target CDSs were aligned with the human genome (*Homo sapiens* GRCh38.p12 chromosomes) by Blastn[17] to obtain matching fragments. Blastn sequence alignment gives an original score of S. To facilitate the comparison of Blast results among different subgenomic groups, the original score is standardized to S’ by Blastn:

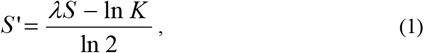

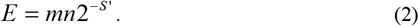

Here, the *E* value represents the expected number of times when two random sequences of length m and n are matched and the score is not lower than *S*’. Parameters *K* and *λ* describe the statistical significance of the results[18]. Assuming that the fragment of length *a* matches perfectly in the two random sequences, one has the following formula:

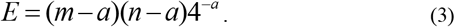

Since the viral genome is quite different from the human genome, matching fragments are usually very short. When *a* is particularly small compared to *m* and *n, a* = *S*’/ 2 is obtained by combining Equation (3) and Equation (4). Thus, HGS is defined as

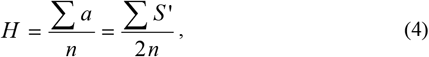

where *n* represents the length of the target sequence. The meaning of *H* is the ratio of the number of matched base pairs to the total length of the sequence when the matched sequences are converted into sequences of the same length.

### Data availability

The SARS-CoV-2 genomes used in this study can be obtained at GISAID website (https://www.gisaid.org/). The SARS-CoV genomes can be obtained at NCBI database (https://www.ncbi.nlm.nih.gov/). The accession number and corresponding HGS of 2594 SARS-CoV-2 genomes and those of 25 SARS-CoV genomes are in Supplemental Information. The code for HGS calculation is available in GitHub (https://github.com/WeitaoNSun/HGS).

## Results

### SARS-CoV-2 ORFs have higher HGS than those of SARS-CoV

The SARS-CoV-2 (GenBank: MN908947.3) and SARS-CoV (GenBank: AY394850.2) RNA sequences were used as references to establish the genome organization. SARS-CoV-2 has 14 5’-ORFs, while SARS-CoV has 19 5’-ORFs. The length of each ORF is no less than 75 nt (**Table 1**).

**Table 1.** Location, length and residue number of each ORF of SARS-CoV-2 and SARS-CoV genomes. The ORF names defined in different papers are listed in the first three columns(9, 18, 19).

| ORF names |  |  | SARS-CoV |  |  |  | SARS-CoV-2 |  |  |  |
| --- | --- | --- | --- | --- | --- | --- | --- | --- | --- | --- |
| Narayanan | Marra | Rota | start | stop | length | residues | start | stop | length | residues |
| ORF1a | ORF1a | 1a | 3361 | 13413 | 10053 | 3350 | 266 | 13483 | 13218 | 4405 |
| N/R | N/R | N/R | 13685 | 13759 | 75 | 24 | 13685 | 13759 | 75 | 24 |
| ORF1b | ORF1b | 1b | 13398 | 21485 | 8088 | 2628 | 13768 | 21555 | 7788 | 2595 |
| S | Sprotein | S | 21492 | 25259 | 3768 | 1282 | 21536 | 25384 | 3849 | 1282 |
| N/R | N/R | N/R | 25207 | 25329 | 123 | 40 | 25332 | 25448 | 117 | 38 |
| ORF3a | ORF3 | X1 | 25268 | 26092 | 825 | 274 | 25393 | 26220 | 828 | 275 |
| E | Eprotein | E | 26117 | 26347 | 231 | 76 | 26245 | 26472 | 228 | 75 |
| M | Mprotein | M | 26398 | 27063 | 666 | 221 | 26523 | 27191 | 669 | 222 |
| ORF6 | ORF7 | X3 | 27074 | 27265 | 192 | 63 | 27202 | 27387 | 186 | 61 |
| ORF7a | ORF8 | X4 | 27273 | 27641 | 369 | 122 | 27394 | 27759 | 366 | 121 |
| ORF7b | ORF9 | N/R | 27638 | 27772 | 135 | 44 | 27756 | 27887 | 132 | 43 |
| ORF8a | ORF10 | N/R | 27779 | 27853 | 75 | 24 | 27894 | 28259 | 366 | 121 |
| ORF8b | ORF11 | X5 | 27862 | 28116 | 255 | 84 | N/R | N/R | N/R | N/R |
| N | Nprotein | N | 28118 | 29386 | 1269 | 422 | 28274 | 29533 | 1260 | 419 |
| N/R | N/R | N/R | 29413 | 29490 | 78 | 25 | 29558 | 29674 | 117 | 38 |

A quantitative definition of HGS was proposed to investigate the similarity between viral coding sequences (CDSs) and the human genome (*Homo sapiens* GRCh38.p12 chromosomes). The CDS alignment scores were determined by using NCBI Blastn[17], and HGS was calculated by the formulas described in the Methods for each ORF in the coronavirus genome. The overall HGS of a full-length virus genome was obtained by the weighted sum of ORF HGSs. The weighting factor was the ratio of ORF length to the full-genome length. The ORF lengths of SARS-CoV and SARS-CoV-2 genomes are given in **Table 1**.

The HGS of ORFs was calculated for 2594 SARS-CoV-2 genomes with geolocation from China, the USA and Europe. Phylogenetic trees representing the HGS relationship among virus strains are shown in Fig 1, Fig 2 and Fig 3 for all three regions. The tree clusters were formed based on the distance between vectors containing ORF HGS values. Most of the genomes had moderate HGS values. Genomes with similar HGS values were usually in the same cluster and shared a common ancestor. The genomes with high HGS were not all concentrated in the same cluster but may form several separate populations in the tree.

**Fig 1.**
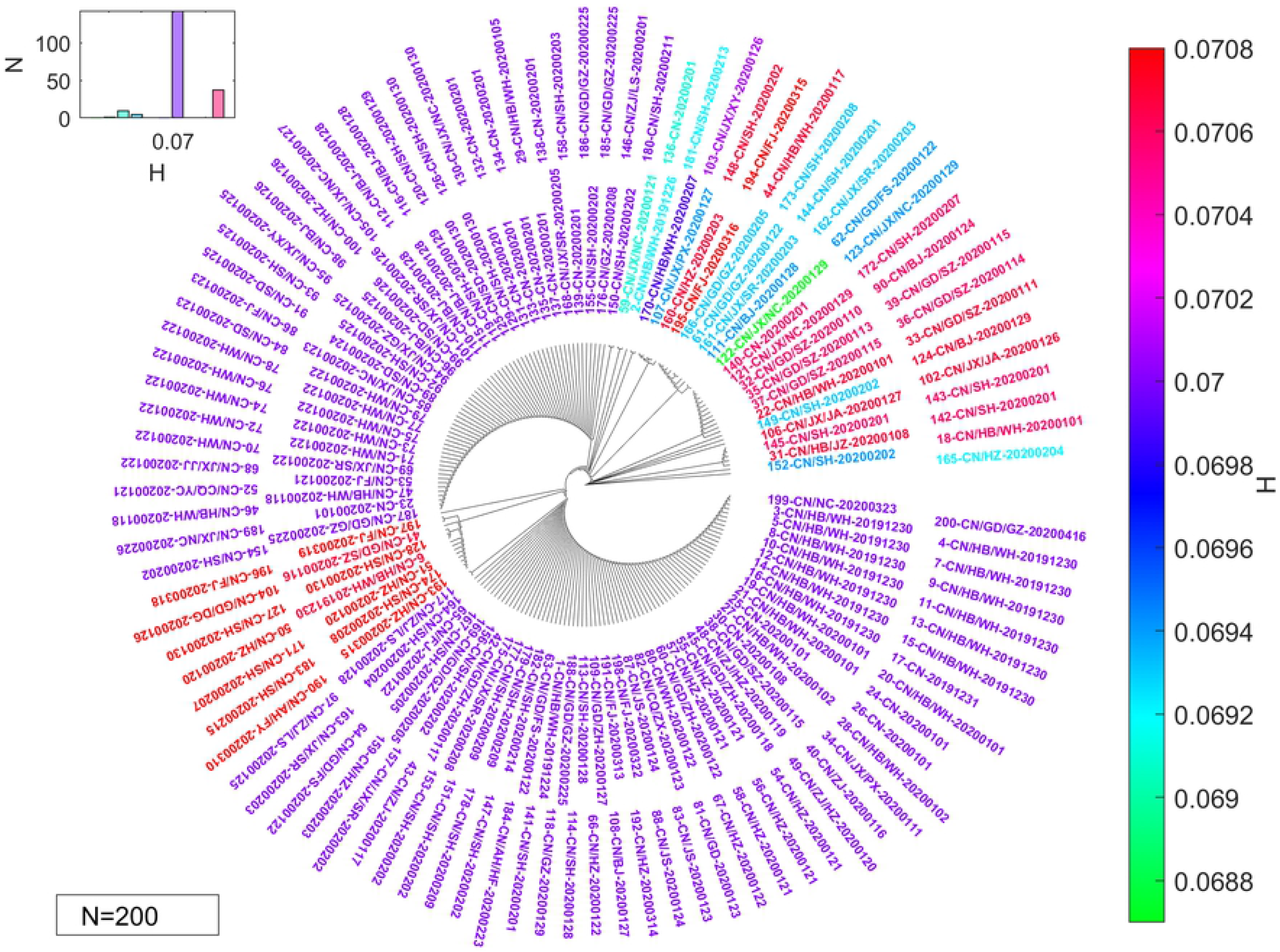
The HGS tree contains 200 SARS-CoV-2 genomes from China. Distance between leaves is the unweighted pair distance between the 10-ORF-HGS vector of genomes. The color bar represents the overall HGS value of each genome (weighted sum of ORF HGS). Out of a total of 200 viral genomes, 36 have unique ORF HGS values. The histogram at the top left shows the distribution of all genome HGS.

**Fig 2.**
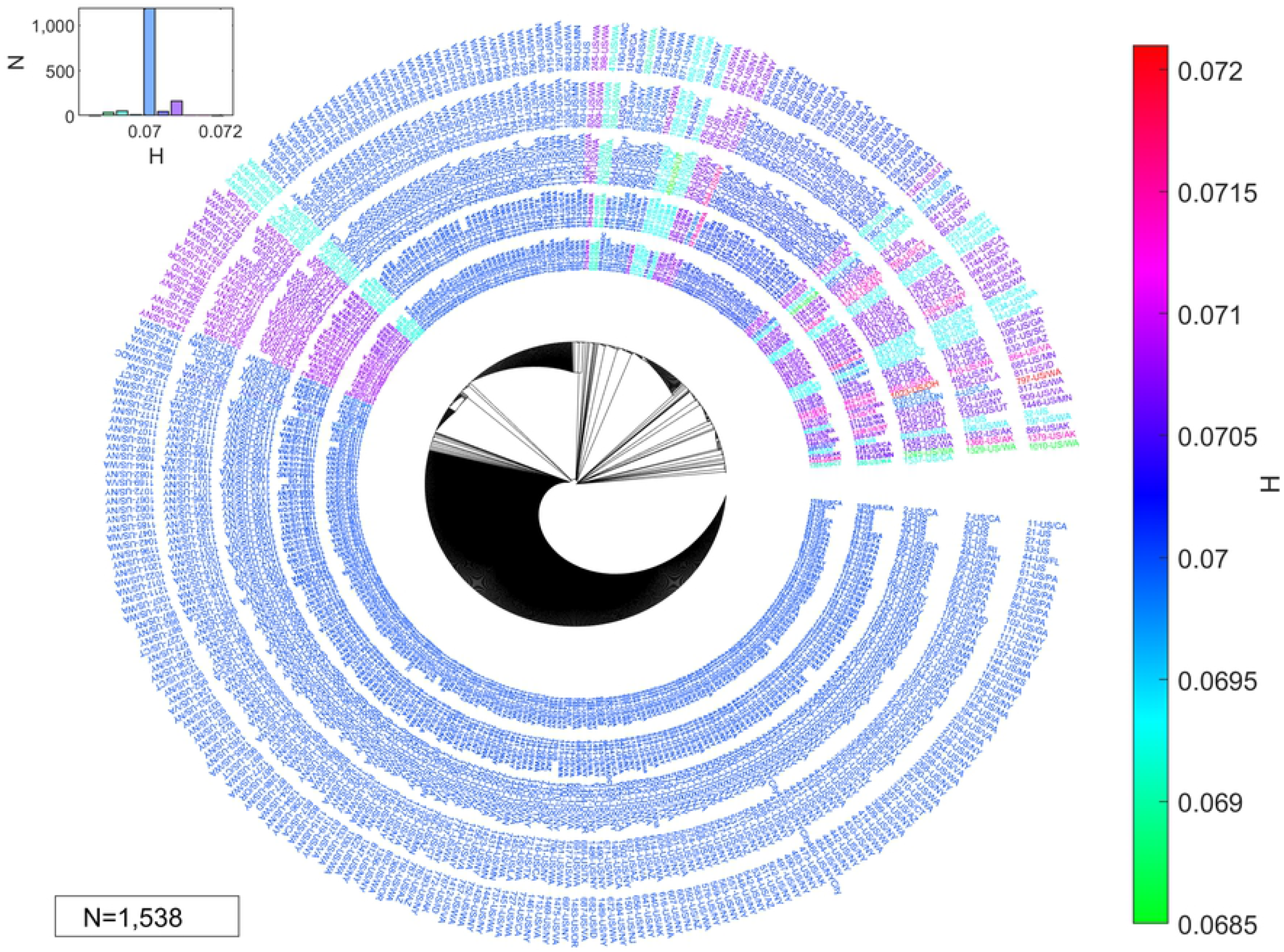
The HGS tree contains 1538 SARS-CoV-2 genomes from the USA. Distance between leaves is the unweighted pair distance between the 10-ORF-HGS vector of genomes. The color bar represents the overall HGS value of each genome (weighted sum of ORF HGS). Out of a total of 1538 viral genomes, 140 have unique ORF HGS values. The histogram at the top left shows the distribution of all genome HGS.

**Fig 3.**
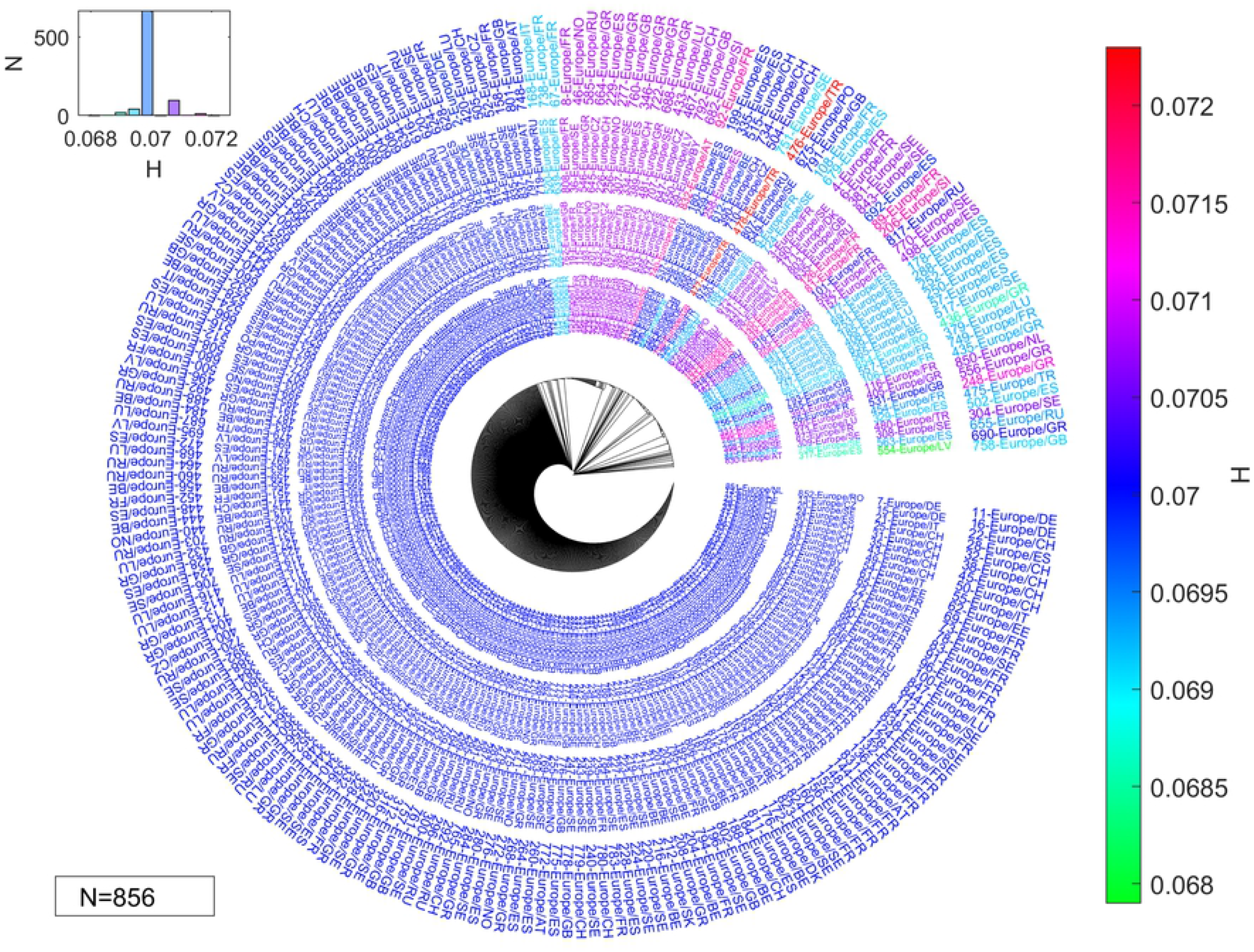
The HGS tree contains 856 SARS-CoV-2 genomes from Europe. Distance between leaves is the unweighted pair distance between the 10-ORF-HGS vector of genomes. The color bar represents the overall HGS value of each genome (weighted sum of ORF HGS). Out of a total of 856 viral genomes, 98 have unique ORF HGS values. The histogram at the top left shows the distribution of all genome HGS.

The full-length genome data were obtained from the Global Initiative on Sharing All Influenza Data (GISAID) database[15]. The sequence requirements were full-length sequences only, sequences with definite collection dates and locations, and no nucleotide names other than A, G, C and T. The number of genomes that met such requirements was 200 for China, 1538 for the USA and 856 for Europe at the time of article preparation. The HGS of human SARS-CoV genomes was also calculated. In NCBI GenBank[16], a total of 25 SARS-CoV CDSs met the above sequence requirements.

**Fig 4** shows that ORF 7b of SARS-CoV had the highest similarity with the human genome, followed by ORF6, ORF7a, ORF3a and ORF 8. For SARS-CoV-2, ORF 7b, ORF 6 and ORF 8 were the top 3 genes with the highest HGSs. The mean HGS values of ORF6 and ORF8 in SARS-CoV-2 increased significantly, reaching 122% and 148% of those of SARS-CoV ORF6 and ORF8, respectively **Fig 4**. The roles of such HGS changes are not clear. However, by investigating the function of the SARS-CoV viral genes and proteins, the mechanism of the rapid spread of the newly emerged COVID-19 may be inferred from the HGS changes in SARS-CoV-2 genomes.

**Fig 4.**
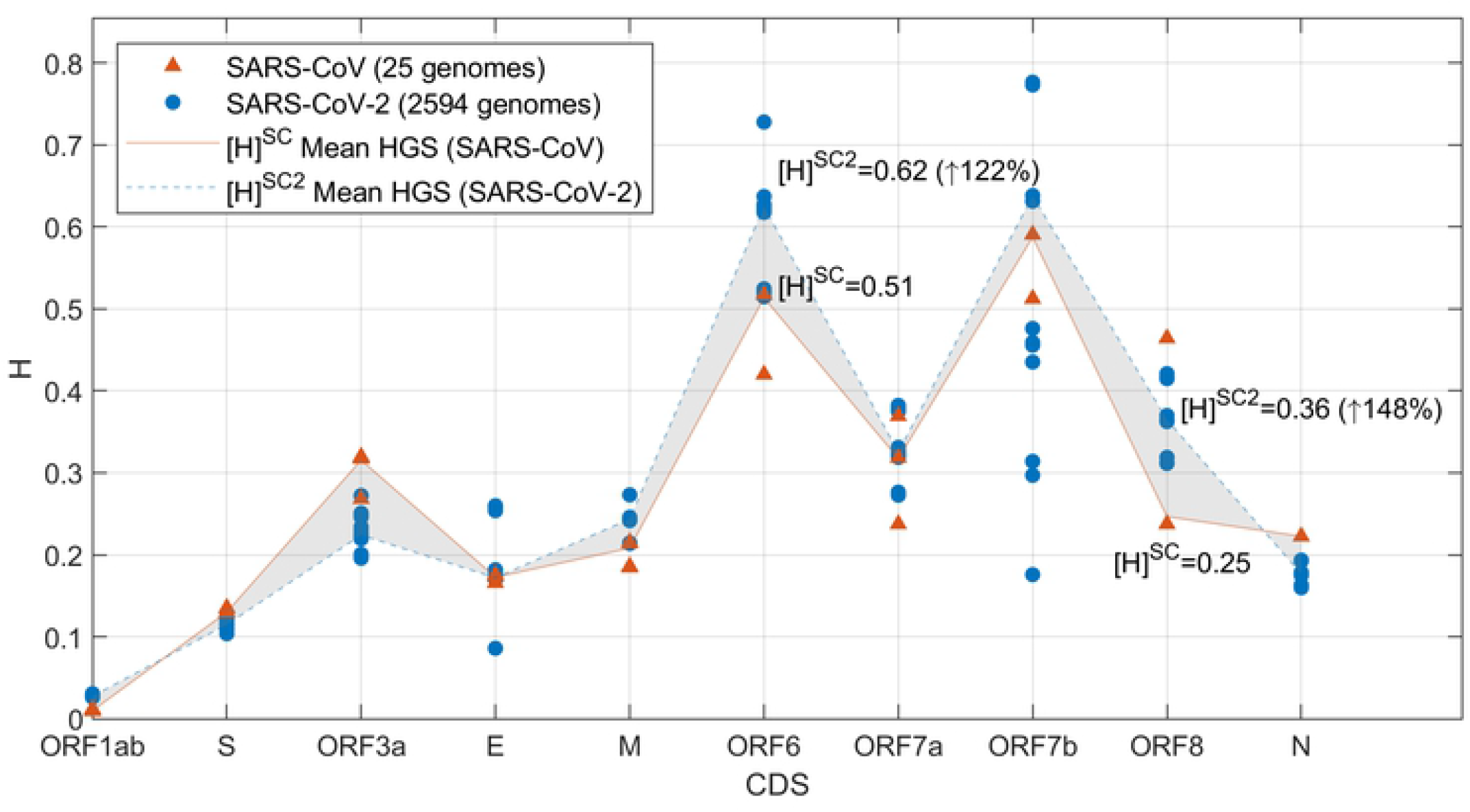
The HGS values of SARS-CoV-2 and SARS-CoV genes. ORF6 and ORF8 of SARS-CoV-2 have apparently higher mean HGS values than those of SARS-CoV, reaching 122% and 148% of that of SARS-CoV ORF6 and ORF8, respectively.

Studies have shown that ORF6 suppresses the induction of IFN and signaling pathways[19]. A membrane protein with 63 amino acids, ORF 6 blocked the IFNAR-STAT signaling pathway by limiting the mobility of the importin subunit KPNB1 and preventing the STAT1 complex from moving into the nucleus for ISRE activation[20]. Laboratory studies confirmed that the expression of ORF 6 transformed a sublethal infection into lethal encephalitis and enhanced the growth of the virus in cells[21]. In addition, ORF 6 circumvented IFN production by inhibiting IRF-3 phosphorylation in the (TRAF3)-(TBK1+IKKε)-(IRF3)-(IFNβ) signaling pathway (**Fig 5**), which is an essential signaling pathway triggered by the viral sensors RIG-1/MDA5 and TLRs[22].

**Fig 5.**
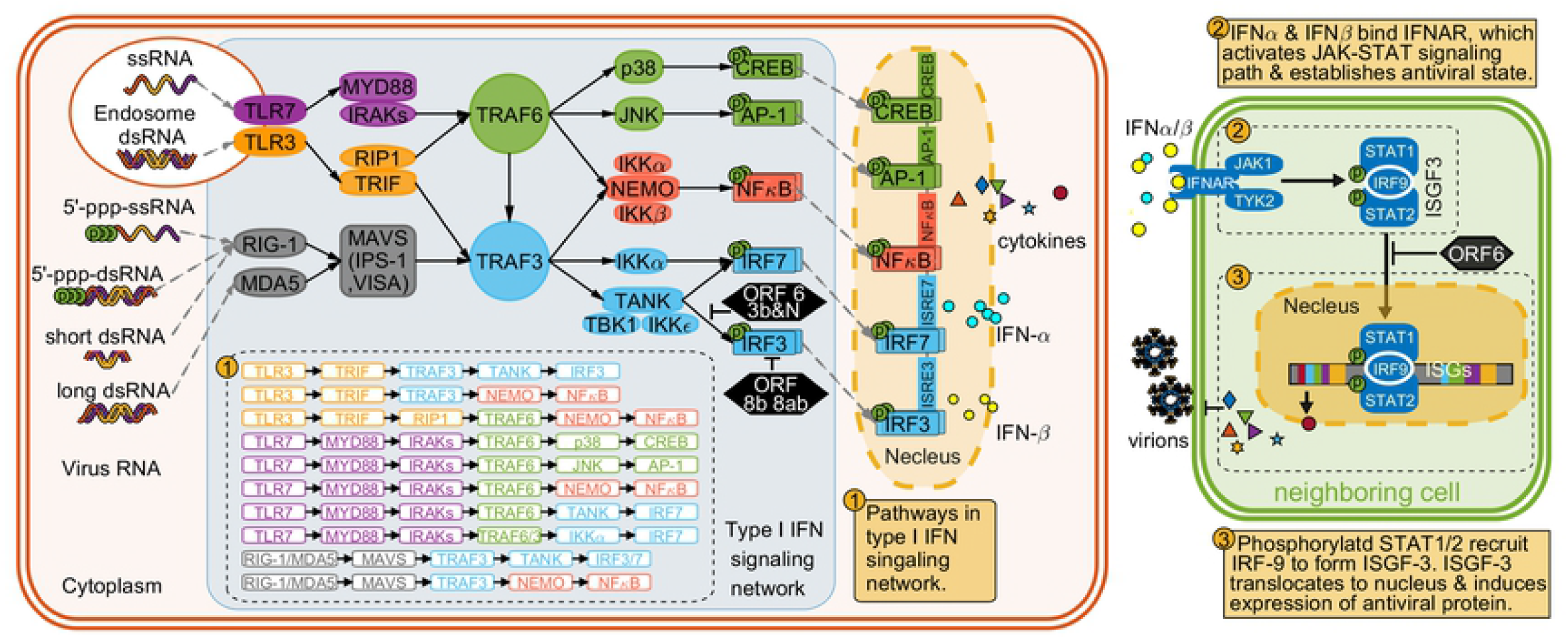
SARS-CoV induced immune response in host cells. Host cell detect virus invasion mainly by TLPs and RIG1/MDA5 and lead to type I IFN signaling pathway. The receptor IFNAR senses type I IFN and leads to the JAK1-STAT signaling pathway, which expresses antiviral proteins and bring neighboring cell into anti-virus state. The ORF6 suppresses type I IFN expression by inhibiting translocation of STAT1+STAT2+IRF9 complex into nucleus. ORF 6 also circumvent IFN production by inhibit IRF-3 phosphorylation in signaling pathway (TRAF3)-(TBK1+IKKε)-(IRF3)-(IFNβ). The expression of ORF8b and 8ab enhance the IRF3 degradation, thus regulating immune functions of IRF3.

An intact gene, ORF8 encodes a single accessory protein at the early stage of SARS-CoV infection and splits into two fragments, ORF8a and ORF8b, at later stages[23]. ORF8a and 8b have been observed in most SARS-CoV-infected cells[24]. Wong et al.[25] found that the proteins ORF8b and ORF8ab in SARS-CoV inhibited the IFN response during viral infection. It was also reported that ORF8b formed insoluble intracellular aggregates and triggered cell death[26]. Amazingly, studies showed that SARS-CoV-related CoVs in horseshoe bats had 95% genome identities to human and civet SARS-CoVs, but the ORF8 protein amino acid similarities varied from 32% to 81%[27]. These findings indicate that the ORF8 gene is more prone than other CoV genes to mutations in virus-host interactions. Overexpression of ORF 8b and ORF 8ab had a significant effect on IRF3 dimerization rather than IRF3 phosphorylation[25]. The 8b region of SARS-CoV protein ORF8 functions in ubiquitination binding, ubiquitination and glycosylation, which may interact with IRF3[28]. The expression of ORF8b and 8ab enhanced IRF3 degradation, thus regulating the immune functions of IRF3 (**Fig 5**). Interestingly, ORF8 is an IFN antagonist expressed in the later stage of SARS-CoV infection. Studies showed that activation of IRF3 was blocked in the late stage of SARS-CoV infection, which was consistent with the late expression of ORF8b. Therefore, the expression of ORF8 may help to suppress the innate immune response that occurs in the later stages of infection and delay IFNβ signaling. This may explain why the virus expresses a late-stage IFN antagonist, such as ORF8.

This work found that genes with high HGS were critical in suppressing innate immunity. Studies have shown that the ORF3b, ORF6 and N proteins of SARS-CoV enhance suppression of IFNβ expression in host innate immunity[29]. When IFN binds to the cell receptor IFNAR, the JAK/STAT signaling pathway is activated, leading to activation of IFN-stimulated genes (ISGs) containing an ISRE in their promoter. Expression of genes with an ISRE will trigger the production of hundreds of antiviral proteins inhibiting viral infections. Therefore, a reduction in expression from ISRE-containing promoters is a direct indicator of the enhanced ability to inhibit IFN synthesis.

ISRE-containing promoter expression after Sendai virus infection needs both IFN synthesis and signaling. However, ISRE-containing promoter expression after IFNβ treatment requires only IFN signaling. In cells treated with IFNβ, it was found that N did not significantly inhibit the expression of the ISRE promoter[29]. The expression level was approximately 78% of the value for the empty control. However, ORF3b and ORF6 still inhibited the expression of the ISRE promoter. We calculated the HGSs of ORF3b, ORF6 and N for SARS-CoV. Amazingly, the results clearly demonstrated that the ISRE-containing promoter expression decreased rapidly with increasing HGS (**Fig 6**), which provided evidence that there was a power law dependence of IFN synthesis inhibition based on HGS. The ISRE-containing promoter expression data followed the work of Kopecky-Bromberg et al.[29]. For 293T cells transfected with the SARS-CoV proteins and infected by Sendai virus[29], IFN inhibition obeys the following power law equation: *P* = 0.004*H* ^−0.539^ + 5.421, where *H* is the HGS value of the viral genes ORF3b, ORF6 and N, and *P* is the expression of genes with an ISRE as a percentage of the value for the empty control. The power law equation for cells treated with IFNβ is *P* = 0.00001*H* ^−11.007^ + 3.633. The coefficient of determination *R*^2^ reaches 1 for both data sets, indicating a perfect fit for the power law dependence on HGS.

**Fig 6.**
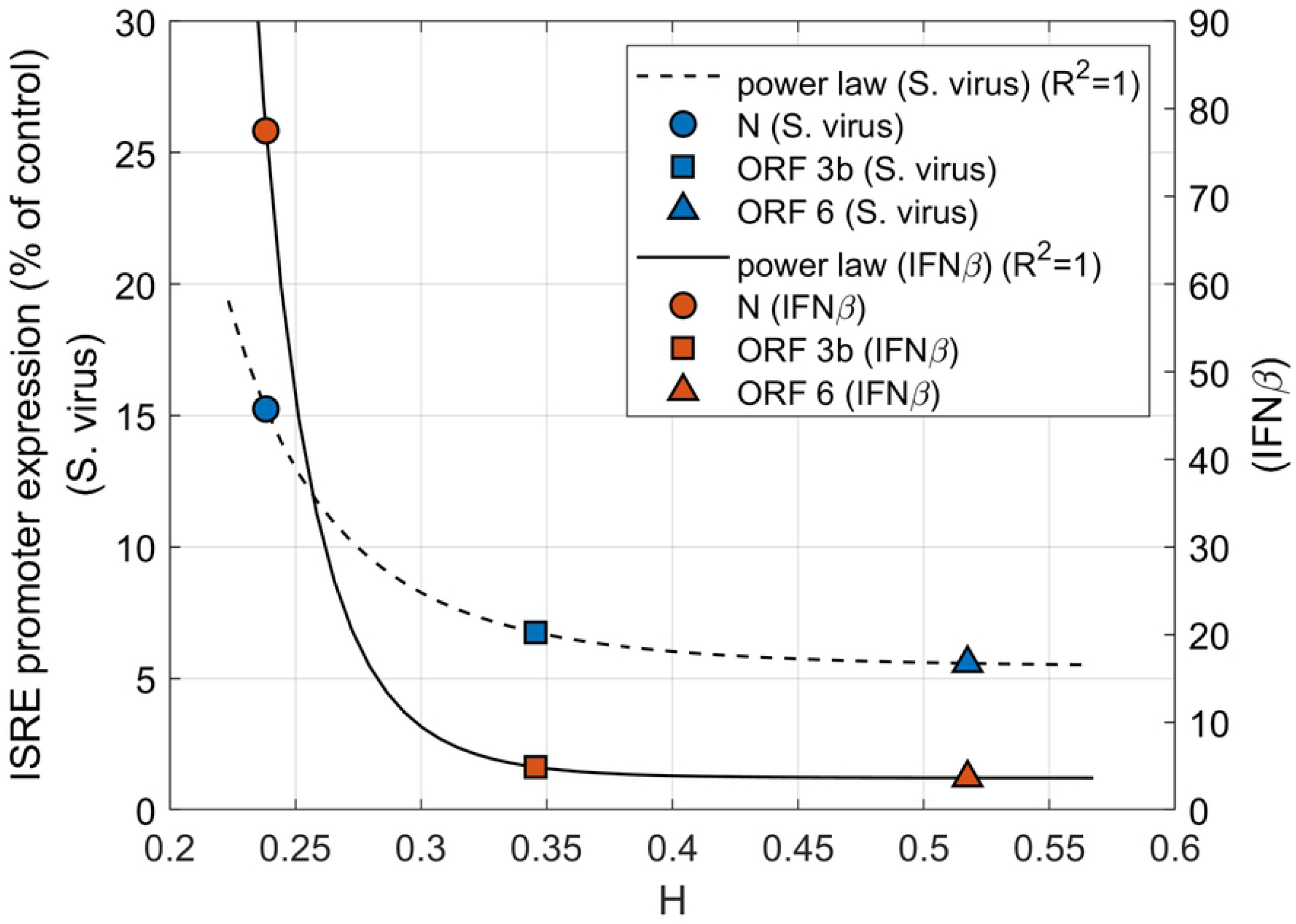
Inhibition of a promoter containing an ISRE by SARS-CoV proteins with different genome HGS values. Cells were cotransfected with the SARS-CoV proteins and either infected with Sendai virus (S. virus) or treated with IFNβ after 24 hours. The expression of the promoter decays rapidly with the increasing HGS of ORF 3b, ORF 6 and N, conforming to a power law.

The findings suggested that HGS, i.e., similarity between the virus and host genome, is a reliable indicator of the suppression of innate immunity by viral proteins. Channappanavar et al. found that rapid SARS-CoV replication and a relative delay in IFN I signaling resulted in immune dysregulation and severe disease in infected mice[30]. Considering the significant HGS increments of ORF6 and ORF8 and their roles in suppressing innate immunity, it could be speculated that SARS-CoV-2 would further suppress IFN I synthesis and delay host innate immunity as HGS increases. This hypothesis may explain the delayed immune response and uncontrolled inflammatory response that lead to the epidemiological manifestations of SARS-CoV-2, such as long incubation periods, mild symptoms, rapid spread and low mortality. However, the mechanism of how viral proteins cause further delay of immune signaling and how it leads to new immunopathological features remain largely unknown.

The discovery of increased HGS of ORF 6 and ORF 8 provides strong evidence that SARS-CoV-2 evolved to be more adapted to humans than SARS-CoV. These inferences offer a valuable picture of how SARS-CoV-2 could have become different from SARS-CoV. In addition, genetic mutations making the virus genome adapted to humans can also be identified through HGS analysis.

### The SARS-CoV-2 mutation 10818G>T is adapted to humans

Recent studies have shown that SARS-CoV-2 had a high mutation rate, and new mutations have emerged in ORF1ab, S, ORF3a and ORF8[31, 32]. However, the types of mutations that contribute to viral adaptations in humans are not clear. To understand how mutations aid survival of SARS-CoV-2 populations under selective pressure, the accumulated nucleotide variants in consensus sequences were identified in 2594 genomes from China, the USA and Europe. The virus genome was identified by its HGS values of ten ORFs (ORF1ab, S, ORF3a, E, M, ORF6, ORF7a, ORF7b, ORF8, and N). The percentages of virus strains with unique ORF HGSs were 18% (36 out of 200), 9% (140 out of 1538) and 11% (98 out of 856) for genomes with geolocations of China, the USA and Europe, respectively. A total of 74 mutations, 162 mutations and 145 mutations were identified in genomes for these three regions, respectively. Gene mutation profiles of SARS-CoV-2 genomes with different HGSs are shown inFig 6, Fig 7 and Fig 8. SARS-CoV-2 in different regions developed its own conserved mutations independently (**Table 2**). For example, the mutations in genomes with a geolocation of China included the ORF1ab mutations 10818G>T (TTG>TTT), 1132G>A (GTA>ATA), and 8517C>T (AGC>AGT); ORF8 mutation 251T>C (TTA>TCA); N mutation 415T>C (TTG>CTG); S mutation 1868A>G (GAT>GGT); and ORF3a mutation 752G>T (GGT>GTT). Here, the number before the mutated nucleotide represents the sequence position relative to the starting point of the ORF where the mutation is located.

**Table 2.** Conserved mutations identified in SARS-CoV-2 genomes with geolocations of China, the USA and Europe.

| mutation | Location(ORF) | protein | residue mutation | residue type | residue property | Geolocation |
| --- | --- | --- | --- | --- | --- | --- |
| TTG>TTT | 10818 | nsp6 | M37F | Phenylalanine | Hydrophobic | China |
| GTA>ATA | 1132 | nsp2 | V198I | Isoleucine | Hydrophobic | China |
| AGC>AGT | 8517 | nsp4 | S76S | Serine | Polar | China |
| TTA>TCA | 251 | ORF8 | L84S | Serine | Polar | China |
| TTG>CTG | 415 | N | M139M | Methionine | Hydrophobic | China |
| GAT>GGT | 1868 | S | D623G | Glycine | Special | China |
| GGT>GTT | 752 | ORF3a | G251V | Valine | Hydrophobic | China |
| ATC>ACC | 794 | nsp2 | I85T | Threonine | Polar | USA |
| CAT>CAG | 171 | ORF3a | H57Q | Glutamine | Polar | USA |
| TTT>TTC | 2772 | nsp3 | F106F | Phenylalanine | Hydrophobic | USA |
| CTT>CCT | 13859 | nsp12 | L228P | Proline | Special | USA |
| GGT>GAT | 1868 | S | G623D | Aspartic acid | Negative | USA |
| TTG>TTT | 10818 | nsp6 | M37F | Phenylalanine | Hydrophobic | USA |
| AGC>AGT | 8517 | nsp4 | S76S | Serine | Polar | USA |
| TTT>TTC | 2772 | nsp3 | F106F | Phenylalanine | Hydrophobic | Europe |
| CTT>CCT | 13859 | nsp12 | L228P | Proline | Special | Europe |
| TTG>TTT | 10818 | nsp6 | M37F | Phenylalanine | Hydrophobic | Europe |
| GGT>GAT | 1868 | S | G623D | Aspartic acid | Negative | Europe |
| GGT>GTT | 752 | ORF3a | G251V | Valine | Hydrophobic | Europe |
| TAC>TAT | 14256 | nsp12 | Y360Y | Tyrosine | Hydrophobic | Europe |
| CAG>CAT | 171 | ORF3a | Q57H | Histidine | Positive | Europe |

**Fig 7.**
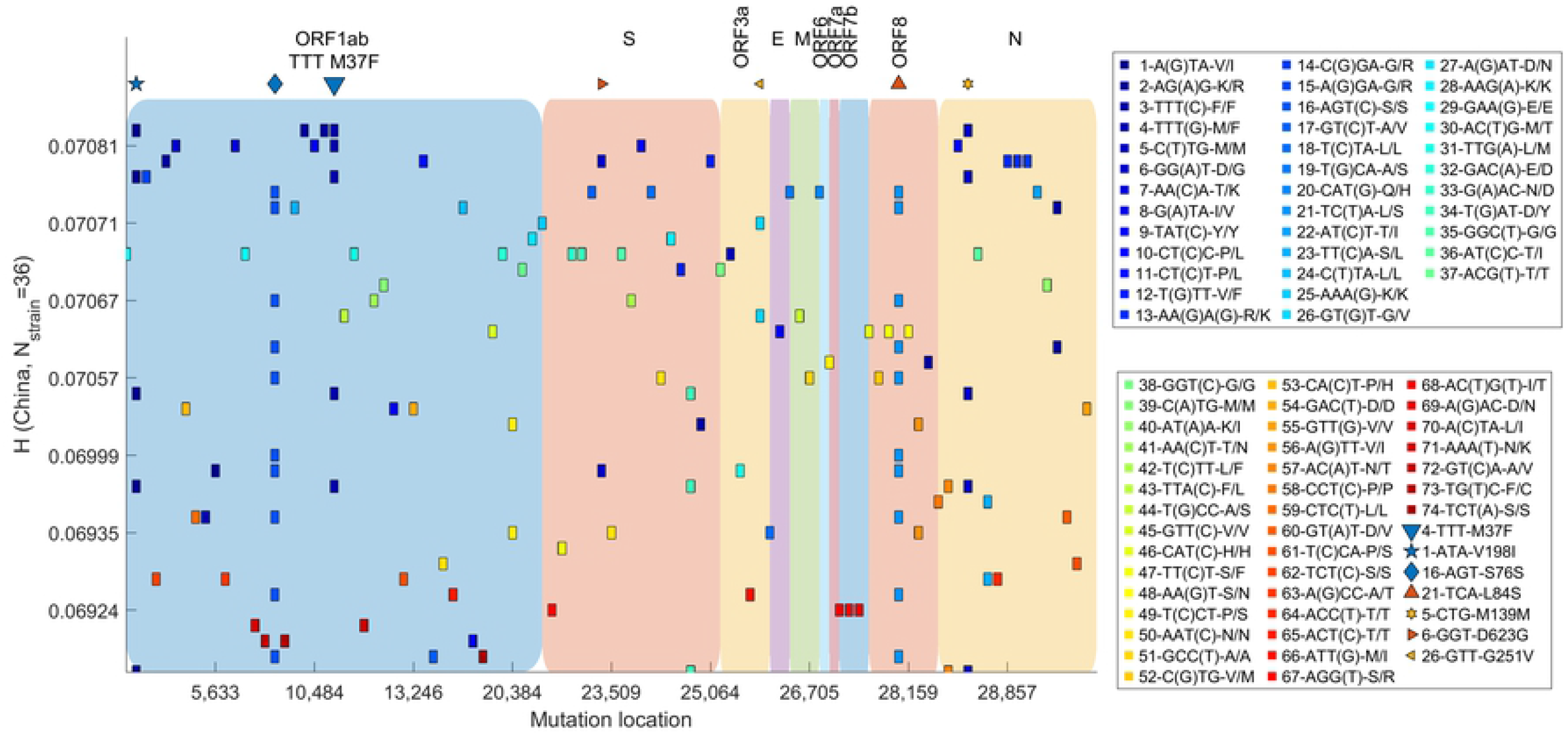
Mutation profile for SARS-CoV-2 genomes (geolocation of China) with different HGS. Out of a total of 200 viral genomes, 36 genomes have unique HGS values. A total of 74 mutations were identified in all the genomes. The top 7 conserved mutations with were shown with special markers at the top of colored blocks representing ORFs. Mutation 10818G>T in ORF1ab (codon TTG>TTT) occurred in populations with high HGS, which results in amino acid M37F mutation in transmembrane protein nsp6. The mutation rarely occurred in populations with low/moderate HGS.

**Fig 8.**
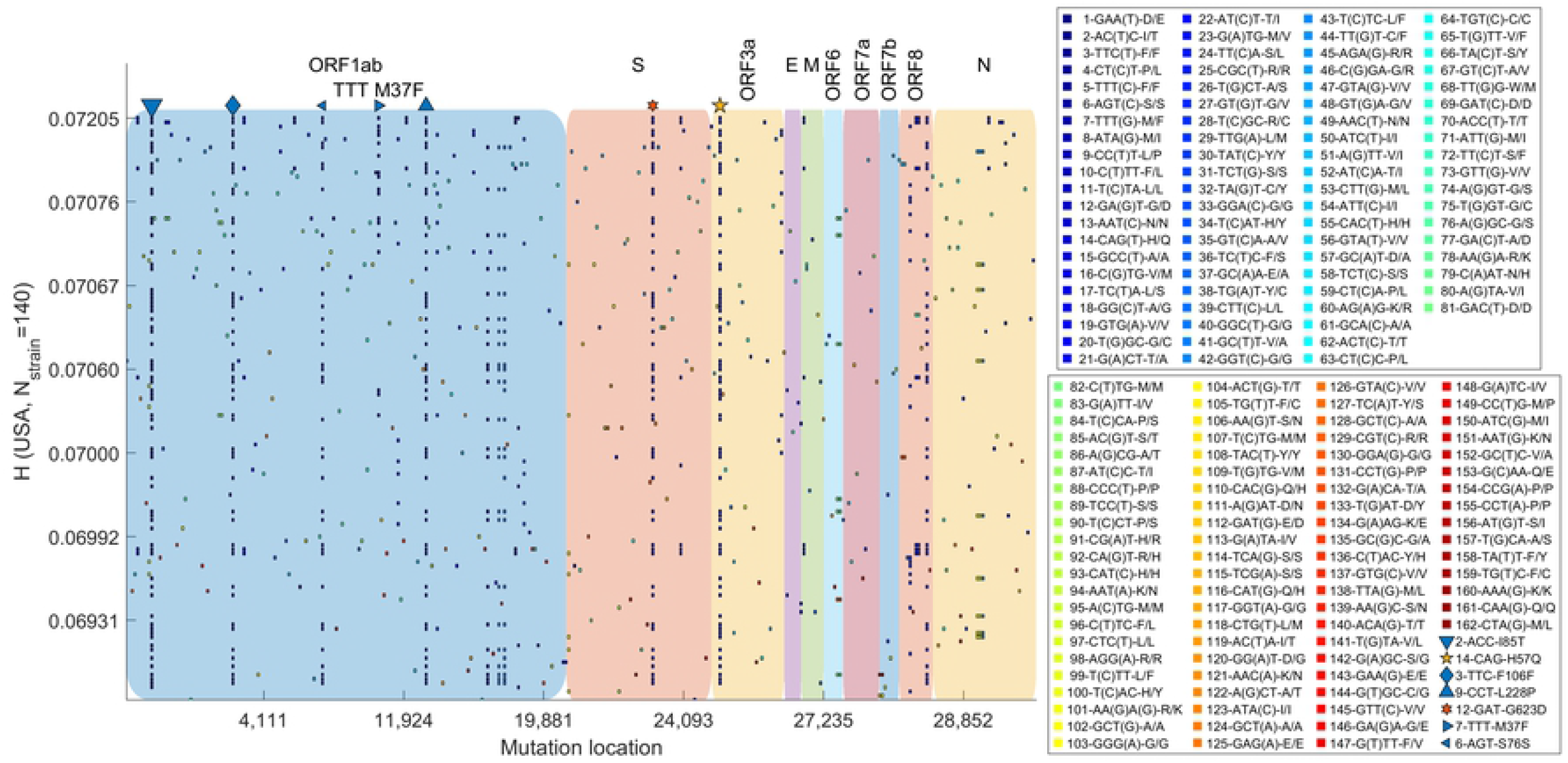
Mutation profile for SARS-CoV-2 genomes (geolocation of the USA) with different HGS. Out of a total of 1538 viral genomes, 140 genomes have unique HGS values. A total of 162 mutations were identified in all the genomes. The top 7 conserved mutations with were shown with special markers at the top of colored blocks representing ORFs. Mutation 10818G>T in ORF1ab (codon TTG>TTT) occurred in populations with high HGS, which results in amino acid M37F mutation in transmembrane protein nsp6. The mutation rarely occurred in populations with low/moderate HGS.

Of all the gene mutations, the ORF1ab 10818G>T(TTG>TTT) mutation is the most interesting. This mutation survived in all three regions (**Fig 10**). In addition, this mutation occurred only in the high HGS population rather than in that with a lower HGS (**Table 3)**. The SARS-CoV-2 ORF1ab gene encodes the precursor polyprotein pp1ab, which is then cleaved into 16 nonstructural proteins (nsp1 to nsp16) by virus-encoded proteinases. nsp6 plays a critical role in membrane anchoring of the RNA replication/transcription complex.

**Table 3.**
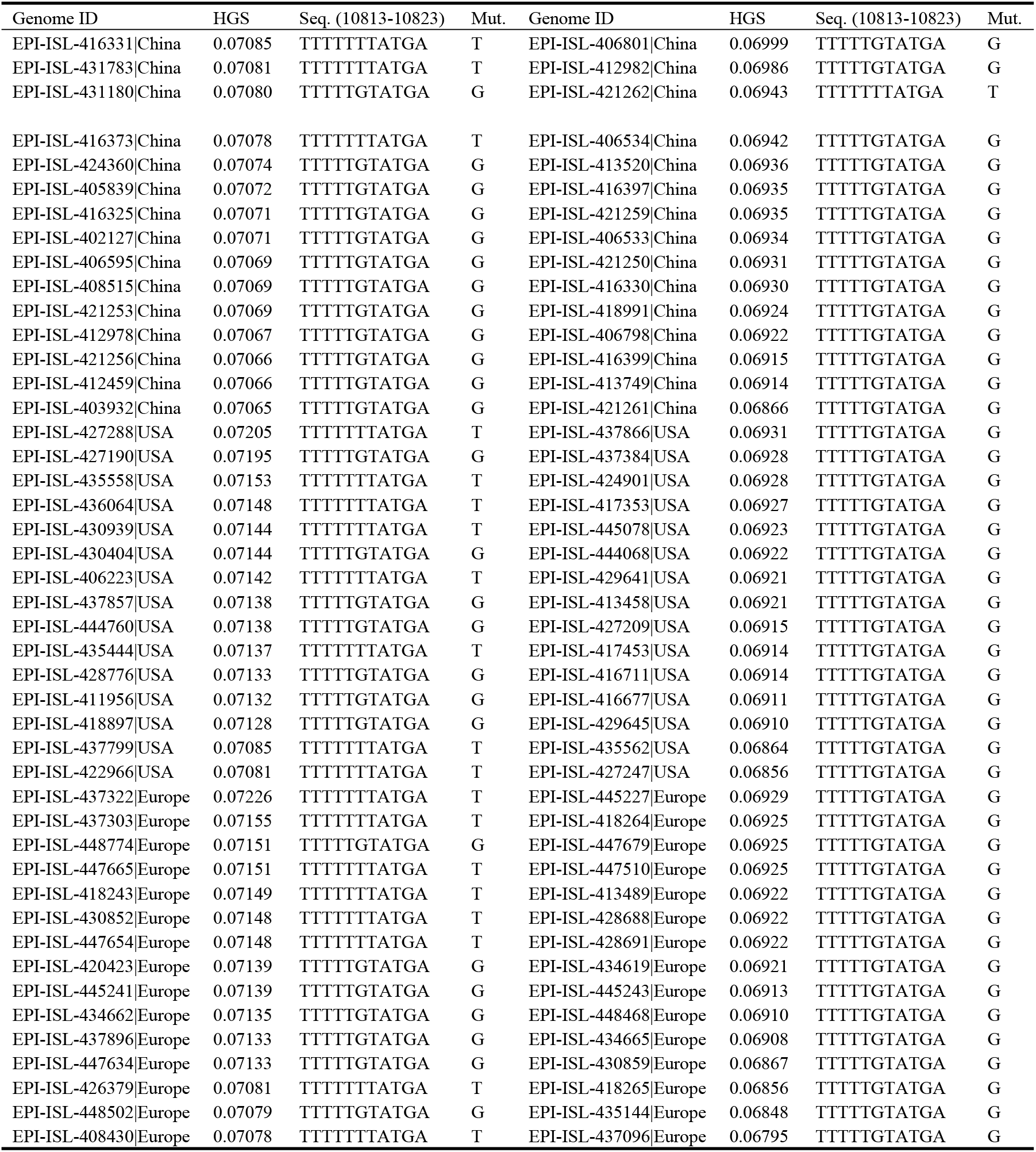
The mutation 10818G>T in ORF1ab (codon TTG>TTT) of SARS-CoV-2 mostly occurs in high-HGS (the first four columns) and rarely occurs in low-HGS population (the last four columns). The top 15 genomes with high HGS are chosen as high-HGS population. The last 15 genomes with low HGS are chosen as low-HGS population. GISAID accession ID and locations are given for genomes from China, the USA and Europe.

**Fig 9.**
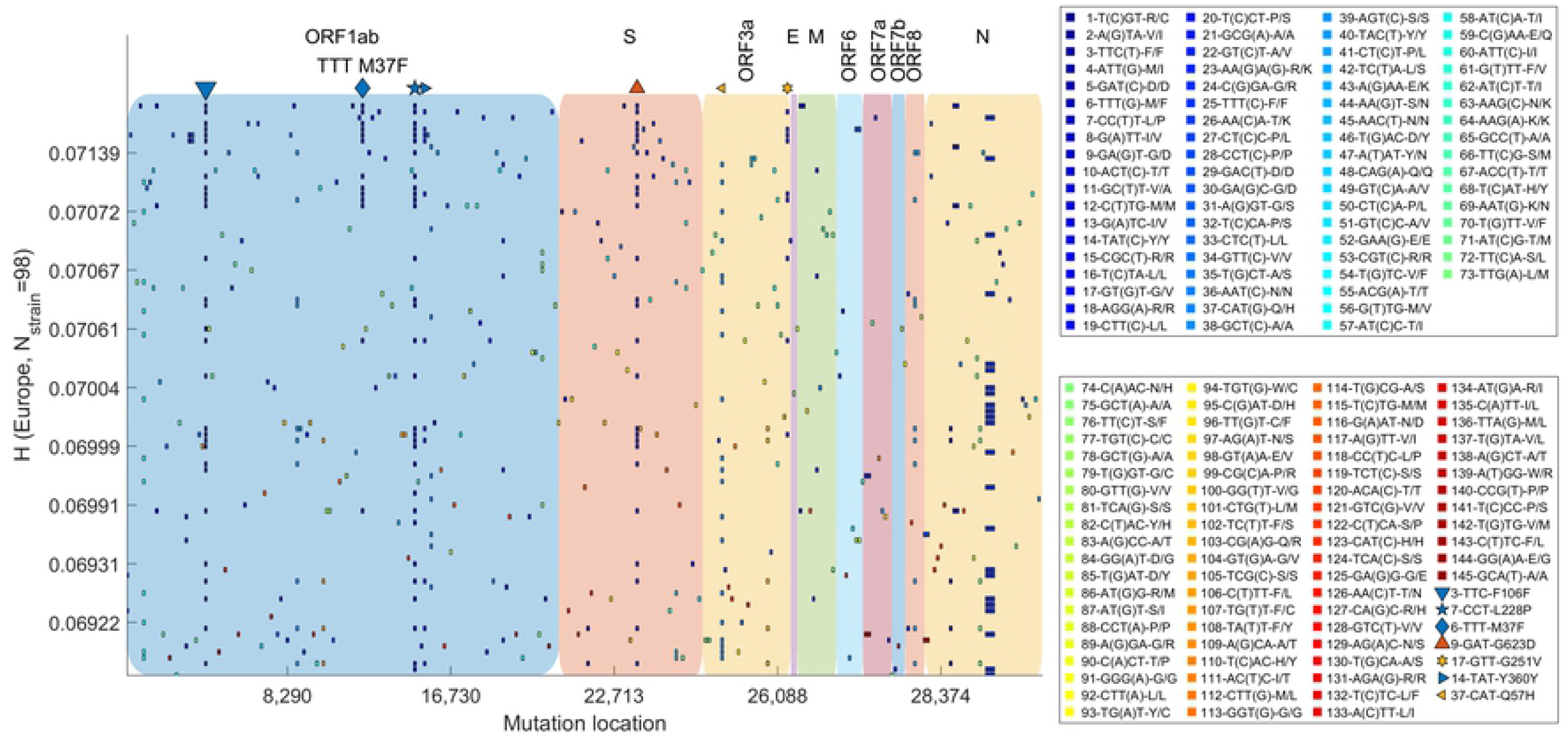
Mutation profile for SARS-CoV-2 genomes (geolocation of Europe) with different HGS. Out of a total of 856 viral genomes, 98 genomes have unique HGS values. A total of 145 mutations were identified in all the genomes. The top 7 conserved mutations with were shown with special markers at the top of colored blocks representing ORFs. Mutation 10818G>T in ORF1ab (codon TTG>TTT) occurred in populations with high HGS, which results in amino acid M37F mutation in transmembrane protein nsp6. The mutation rarely occurred in populations with low/moderate HGS.

**Fig 10.**
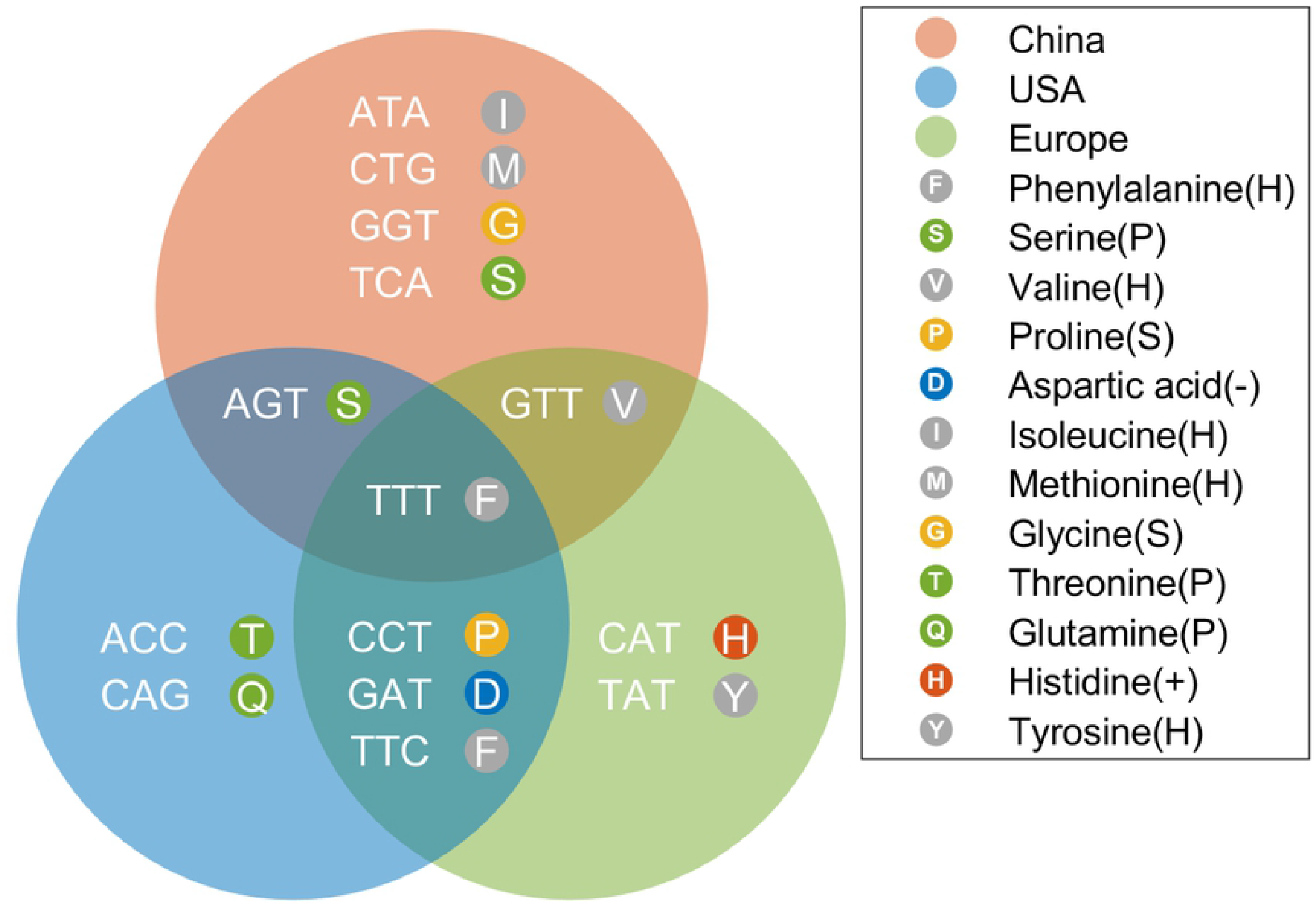
Highly conserved mutations identified in SARS-CoV-2 genomes with geolocations of China, the USA and Europe. The three regions have different sets of mutations. The TTT (F, Phenylalanine) mutation occurred in all three regions. TTT represents the mutation 10818G>T(TTG>TTT) in ORF1ab. The F in the circle represents the amino acid mutation M37F (Methionine to Phenylalanine) in nonstructural protein nsp6. The P, H, +, - and S in brackets in the legend represent polar, hydrophobic, positively charged, negatively charged and special residues, respectively.

The expression of the nonstructural protein nsp6 along with nsp3 and nsp4 mediates the formation of double-membrane vesicles (DMVs)[33], which are organelle-like structures for viral genome replication and protect against host cell defenses.

Studies on the nsp6 protein showed that the protein is a transmembrane protein with 6 transmembrane regions[34]. This 10818G>T ORF1ab mutation caused an amino acid mutation, M37F, in the nonstructural protein nsp6, which is located in a loop between the first and second transmembrane domains on the N-terminal side (**Fig 11**). This finding strongly suggested that the 10818G>T (M37F) mutation survived a selection event and resulted in a new population of SARS-CoV-2 with high HGS, which could be more adapted to humans. In addition, the simultaneous occurrence of ORF1ab 10818G>T in all three regions demonstrated that the mutation was highly stable in human-adapted strains. Although mutations in the nonstructural proteins nsp4 and nsp6 may affect the assembly of DMVs and viral autophagy, the underlying basis of how the M37F mutation results in SARS-CoV-2 adaptation in humans is not clear.

**Fig 11.**
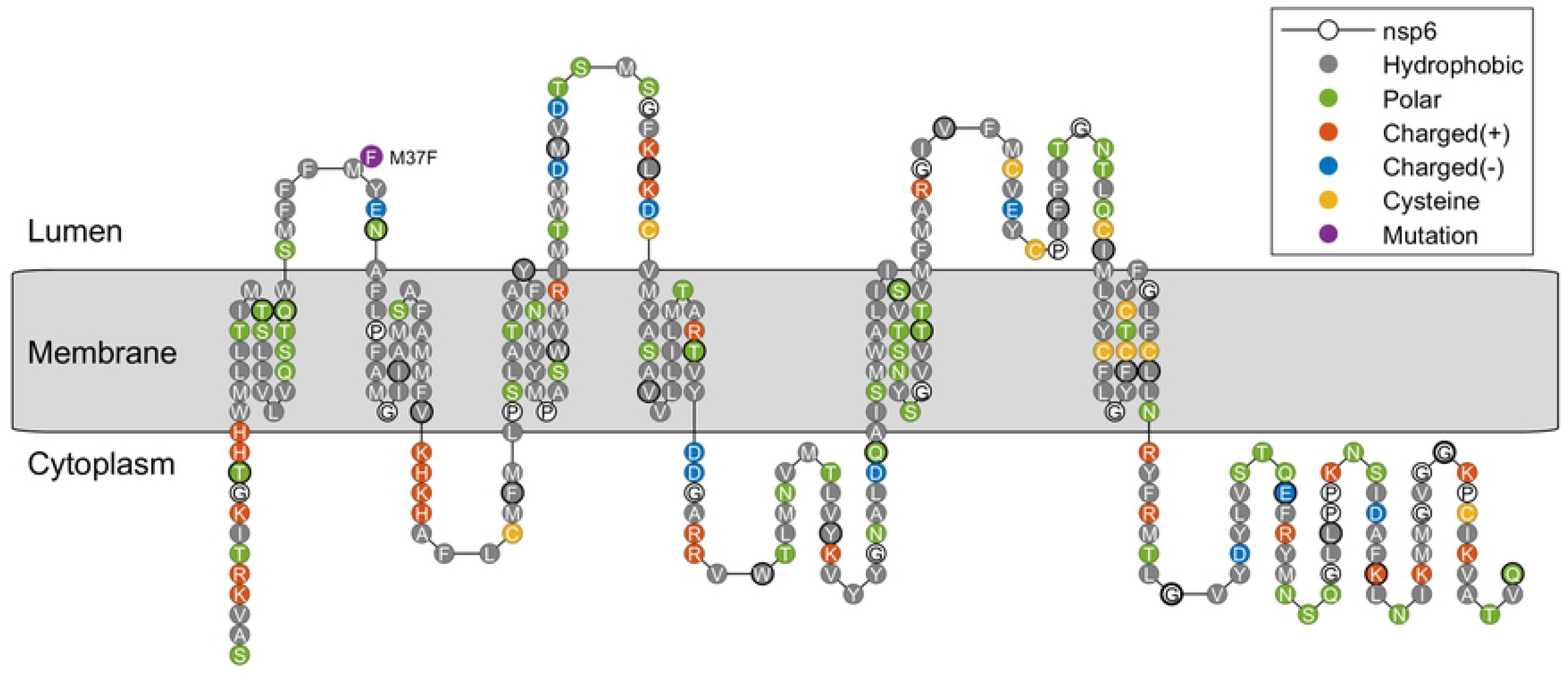
The topology of transmembrane protein nsp6 and the identified M37F mutation located in a loop between the first and second transmembrane domains on the N-terminal side.

The identification of conserved mutations demonstrates that SARS-CoV-2 can improve host adaptation. It is reasonable to hypothesize that high HGS in SARS-CoV-2 genomes and conserved mutations may explain the epidemiological characteristics of COVID-19, such as mild symptoms, rapid spread and low mortality. However, the mechanism behind the impairment remains poorly understood and calls for future laboratory investigations. Viral genome data

## Discussion

The HGS differences between SARS-CoV-2 and SARS-CoV genomes are critical to understanding clinical manifestations of the ongoing pandemic. ORF 6, 7b, 8 are the top 3 genes with significant HGS in SARS-CoV-2. What’s more, ORF 6 and ORF 8 of the SARS-CoV-2 have clear increments in HGS, up to about 122% and 148% of that of SARS-CoV. Such apparent HGS changes suggest that these ORFs are important in defining the difference between SARS-CoV-2 and SARS-CoV. In the ongoing SARS-CoV-2 pandemic, the number of infected people is growing much faster than SARS and the total number of diagnosed cases exceeded that of SARS. But the mortality rate (about 3 %) was lower than SARS (about 11%)[35]. It is known that the primary targets of SARS-CoV are lung and small intestine[36, 37]. Recent studies have found that the SARS-CoV-2 may impair kidney function[38], infect the digestive system[39] and heart[40], and cause liver damage[41]. Recent study showed that SARS-CoV-2 can cause thromboembolic complications[42]. It has been reported that the SARS-CoV-2 virus can be found in stools and urine[40, 43]. In addition, an unusually long incubation period has been reported, during which more than half of the patients had no signs of disease and the virus carriers may be highly contagious [43]. Why the COVID-19 is so different from SARS is still not clear. But the mutations in virus genome and encoded proteins (such as spike protein S) are believed as an important factor.

The knowledge on SARS-CoV-2 accessory proteins by now is quite limited. However, the viral genome and proteins of SARS-CoV have been studied in depth in the past decade. Coronavirus has evolved to escape the innate immune (especially IFN-I expression and signaling) through suppression of IFN induction and singling pathways by non-structural proteins (nsps), structural proteins (S, E, M, N), and accessory proteins (ORF 3a, 6, 7a, 7b, 8a, 8b) [20, 44-53]. By comparing the SARS-CoV gene HGS with that of SARS-CoV-2, the obvious host-genome similarity changes shed light on the cause of rapid spread of COVID-19.

A power-law relationship is recognized between HGS and the expression of ISRE promoter, which is a direct indicator of the virus to inhibit interferon synthesis. The HGS of ORF6 and ORF8 increase greatly in SARS-CoV-2, which represents enhanced ability in suppressing innate immune. Although the functions of accessory proteins of SARS-CoV-2 have not been well studied, the secondary structure prediction reveals that ORF 6 and 8 are transmembrane proteins and may have related functions as in SARS-CoV. In fact, the SARS-CoV-2 contains a full-length ORF 8, which in SARS-CoV this reading frame is divided into ORF 8a and ORF 8b. Linking of ORF 8a and ORF 8b into a single continuous gene fragment had no significant effect on virus growth and RNA replication *in vitro*[54], which indicates that there are ORFs of SARS-CoV-2 may be similar to ORFs of SARS-CoV in function.

The discovery of increased HGS of ORF 6 and ORF 8 provide a strong evidence that SARS-COV-2 evolved to be more adaptable to humans than SARS-CoV. Based on these findings, following conjecture is proposed that the SARS-CoV-2 genes involved in suppressing the host’s innate immunity are more powerful. Therefore, SARS-CoV-2 causes the delayed response of host innate immunity, which results in rapid transmission, low mortality and asymptomatic infection. These inferences are based on bioinformatics data, but offer a valuable picture of how SARS-CoV-2 could become different from SARS-CoV. In addition, the HGS method can also identify genetic mutations that help the virus adapt to humans.

It took the coronavirus 17 years to update from SARS-CoV to SARS-CoV-2. The significant increase in host-genome similarity distinguishes SARS-COV-2 from SARS-COV. SARS-CoV-2 found out a way to improve host adaptation. It is reasonable that high HGS may explain the quite different epidemiological characteristics of SARS-CoV-2, such as mild symptoms, rapid spread and low mortality. But the mechanism behind the impairment remains poorly understood and calls for future laboratory investigations. The COVID-19 appears to be less able to cause deaths than SARS and MERS during the ongoing pandemic. However, there is still a serious warning sign about viral mutation. The threat of another coronavirus outbreak with high infectiousness and mortality remains an alarming possibility.

## Funding

This work was supported by the C.C. Lin specific fund.

## Acknowledgments

I thank the laboratories for sharing SARS-CoV-2 and SARS-CoV genomes through GISAID and NCBI database.

## Supporting information

**Dataset S1 (separate file)**. The accession number and corresponding HGS of 200 SARS-CoV-2 genomes with geolocation of China. Filename is DatasetS1_China_SARS-CoV-2_nstrain200_ORFHGS_allinone.xls. The file contains accession ID, collection date, location, HGS values for 10 ORFs (ORF1ab, S, ORF3a, E, M, ORF6, ORF7a, ORF7b, ORF8, and N) and the weighted HGS of the whole genome.

**Dataset S2 (separate file)**. The accession number and corresponding HGS of 1538 SARS-CoV-2 genomes with geolocation of the USA. Filename is DatasetS2_USA_SARS-CoV-2_nstrain1538_ORFHGS_allinone.xls. The file contains accession ID, collection date, location, HGS values for 10 ORFs (ORF1ab, S, ORF3a, E, M, ORF6, ORF7a, ORF7b, ORF8, and N) and the weighted HGS of the whole genome.

**Dataset S3 (separate file)**. The accession number and corresponding HGS of 856 SARS-CoV-2 genomes with geolocation of Europe. Filename is DatasetS3_Europe_SARS-CoV-2_nstrain856_ORFHGS_allinone.xls. The file contains accession ID, collection date, location, HGS values for 10 ORFs (ORF1ab, S, ORF3a, E, M, ORF6, ORF7a, ORF7b, ORF8, and N) and the weighted HGS of the whole genome.

**Dataset S4 (separate file)**. The accession number and corresponding HGS of 25 SARS-CoV genomes. Filename is DatasetS4_SARS-CoV_nstrain25_ORFHGS_allinone.xls. The file contains accession ID, HGS values for 10 ORFs (ORF1ab, S, ORF3a, E, M, ORF6, ORF7a, ORF7b, ORF8, and N) and the weighted HGS of the whole genome.

